## Supplementary material for "Accessibility of the unstructured α-tubulin C-terminal tail is controlled by microtubule lattice conformation": Key Resources Table (from another version of this preprint: version 2, 2025-12-19)

| REAGENT or RESOURCE | SOURCE | IDENTIFIER |
|---|---|---|
| **Antibodies** | | |
| anti-β-tubulin, mouse monoclonal, clone E7 (1:1,000 immunofluorescence) | DSHB | Cat# E7; RRID: AB_528499 |
| Anti-PA tag, rat monoclonal, clone NZ-1 (1:1,000 western blotting) (1:500 immunofluorescence) | FUJIFILM Wako Pure Chemicals | Cat# 016-25861 |
| Anti-detyrosinated α-tubulin, rabbit monoclonal, clone RM444 (0.1 µg/ml western blotting)(0.5 µg/ml immunofluorescence) | RevMAb Biosciences | Cat# 31-1335-00 |
| Anti-tyrosinated α-tubulin, rat monoclonal, clone YL1/2 (1:1,000 western blotting) | Bio-Rad | Cat# MCA77G |
| Anti-α-tubulin, mouse monoclonal, clone DM1α (1:3,000 western blotting) | MilliporeSigma | Cat# 05-829 |
| anti-GFP, chicken polyclonal (1:1,000 immunofluorescence) | Aves labs | Cat# GFP-1010 |
| Anti-GST, mouse monoclonal (1:1,000 western blotting) | Nacalai USA | Cat# 04435-26 |
| anti-GAPDH, mouse monoclonal, clone G-9 (1:2,000 western blotting) | Santa Cruz | Cat# sc-365062 |
| anti-TagRFP, mouse monoclonal, clone 6A11f (1:1,000 western blotting) | Kerafast | Cat# EFH005 |
| anti-RFP (mScarlet), rabbit polyclonal, (1:1,000 western blotting) | Rockland Immunochemicals | Cat# 600-401-379 |
| Anti-mouse IgG Alexa Fluor 680 AffiniPure, donkey polyclonal (1:500 immunofluorescence) | Jackson ImmunoResearch | Cat# 715-625-150 |
| anti-rabbit IgG Alexa Fluor 594, goat polyclonal (1:1,000 immunofluorescence) | Thermo Fisher | Cat# A-11012 |
| anti-rat IgG Alexa Fluor 488 (1:1,000 immunofluorescence) | Thermo Fisher | Cat# A-11006 |
| anti-chicken IgY Alexa Fluor 488 (1:1,000 immunofluorescence) | Thermo Fisher | Cat# A-11039 |
| Anti-α-tubulin Alexa Fluor 647, mouse monoclonal, clone DM1α (1:500 immunofluorescence) | MilliporeSigma | Cat# 05-829-AF647 |
| Anti-rat IgG Alexa Fluor 680, goat (1:5,000 western blotting) | Thermo Fisher | Cat# A-21096 |
| Anti-mouse IgG Alexa Fluor 700, goat (1:5,000 western blotting) | Thermo Fisher | Cat# A-21036 |
| Anti-mouse IgG DyLight 800, goat (1:10,000 western blotting) | Thermo Fisher | Cat# SA5-10176 |
| Anti-rat IgG DyLight 800, donkey (1:5,000 western blotting) | Thermo Fisher | Cat# SA5-10032 |
| Anti-rabbit IgG IRDye 800CW, goat (1:10,000 western blotting) | LI-COR | Cat# 926-32211 |
| **Chemicals and biological reagents** | | |
| Biotinylated-porcine brain tubulin | Cytoskeleton | Cat# T333P |
| HiLyte647 porcine brain tubulin | Cytoskeleton | Cat# TL670M |
| HiLyte488 porcine brain tubulin | Cytoskeleton | Cat# TL488M |
| Bovine brain tubulin | R. Ohi lab | N/A |
| HeLa S3 tubulin | Ohi and Verhey labs | Thomas et al. 2025 |
| Alexa Fluor 647 HeLa S3 tubulin | Ohi and Verhey labs | Thomas et al. 2025 |
| Alexa Fluor 647 HeLa S3 tubulin | Ohi and Verhey labs | Thomas et al. 2025 |
| Taxol | Cytoskeleton | Cat# TXD01 |
| puromycin | MilliporeSigma | Cat# P8833 |
| doxycycline | Thermo Fisher | Cat# BP26531 |
| Lipofectamine 2000 | Thermo Fisher | Cat# 11668019 |

| | | |
|---|---|---|
| Opti-MEM | Thermo Fisher | Cat# 31985070 |
| Bovine serum albumin | MilliporeSigma | Cat# A9647 |
| Casein | MilliporeSigma | Cat# C8654 |
| Glycerol | Thermo Fisher | Cat# BP229-4 |
| Glucose | MilliporeSigma | Cat# G7528 |
| Glucose oxidase | MilliporeSigma | Cat# G7141-10KU |
| Catalase | MilliporeSigma | Cat# C3515 |
| GTP | MilliporeSigma | Cat# G8877 |
| GMPCPP | Jena Bioscience | Cat# NU405S |
| ADP | MilliporeSigma | Cat# A2754 |
| hexokinase | MilliporeSigma | Cat# H5000 |
| apyrase | MilliporeSigma | Cat# A6535 |
| ATP | MilliporeSigma | Cat# A7699 |
| Janelia Fluor X 554 (JFX554) Halo ligand | Janelia Farms | Cat# JFX554 |
| BSA-biotin | MilliporeSigma | Cat# A8549 |
| NeutrAvidin | Thermo Fisher | Cat# 31000 |
| fish skin gelatin (FSG) | MilliporeSigma | Cat# G7765 |
| Lysozyme | MilliporeSigma | Cat# L6876 |
| SIGMAFAST protease inhibitor cocktail | MilliporeSigma | Cat# S8830 |
| Benzonase nuclease | MilliporeSigma | Cat# E1014 |
| Biotin | MilliporeSigma | Cat# B4501 |
| **Bacterial strains** | | |
| DH5α | Thermo Fisher | Cat# 18258-012 |
| Rosetta2(DE3)pLysS | Novagen | Cat# 71403-3 |
| BL21-CodonPlus-RILC | Aligent Technologies | Cat# 230245 |
| DH10Bac | Thermo Fisher | Cat# 10361012 |
| **Cell lines and media** | | |
| COS-7 cells, male *Ceropithecus aethiops* (green monkey) kidney fibroblast | ATCC | RRID: CVCL_0224 |
| Dulbecco's modified Eagle medium (DMEM) | Gibco, Thermo Fisher | Cat# 11960044 |
| Fetal Clone III | HyClone | Cat# SH3010903 |
| GlutaMAX (L-alanyl-L-glutamine dipeptide in 0.85% NaCl) | Gibco, Thermo Fisher | Cat# 35050061 |
| HeLa Kyoto cells, female *Homo sapiens* | Shuh Narumiya | RRID: CVCL_1922 |
| Knock-in HeLa Kyoto cell lines co-expressing 4xCAPGly-mEGFP, 4xCAPGly-mSc3 and sTagRFP-A1aY1 | This study | N/A |
| Dulbecco's modified Eagle medium (DMEM) | Gibco, Thermo Fisher | Cat# 11965118 |
| Fetal Bovine Serum (FBS) | Cytiva | Cat# SH3007103T |
| Penicillin-Streptomycin | Gibco, Thermo Fisher | Cat# 15140122 |
| HeLa S3 cells, female *Homo sapiens* | ATCC (CCL-2.2) | RRID:CVCL_0058 |
| Dulbecco's modified Eagle medium (DMEM) | Gibco, Thermo Fisher | Cat# 11965092 |
| Leibovitz's L-15 Medium | Gibco, Thermo Fisher | Cat# 21083027 |
| Sf9 cells | Thermo Fisher | Cat# 11496015 |
| sf900 II SFM medium | Thermo Fisher | Cat# 10902088 |
| antibiotic antimycotic | Gibco, Thermo Fisher | Cat# 15240062 |
| Cellfectin II | Thermo Fisher | Cat# 10362100 |
| Grace's insect cell culture medium | Gibco, Thermo Fisher | Cat# 11595-030 |

| | | |
|---|---|---|
| **Other** | | |
| poly-D-lysine coated glass-bottom dishes | MatTek | Cat# P35GC-1.5-14-C |
| Prolong Gold | Thermo Fisher | Cat# P36930 |
| Prolong Diamond | Thermo Fisher | Cat# P36970 |
| HiLoad 16/600 Superdex 200 prep grade column | Cytiva | Cat# 28989335 |
| Strep-Tactin XT 4Flow resin | Iba Life Sciences | Cat# 2-5010-002 |
| Fluorescent Protein Labeling Kit | Thermo Fisher | Cat# A10235 |
| Fab Preparation Kit | Thermo Fisher | Cat# 44985 |
| HiPure Plasmid DNA miniprep kit | Thermo Fisher | Cat# K20003 |
| **Recombinant DNA** | | |
| pN1-4xCAPGly-mEGFP | This study | Lab plasmid |
| pN1-4xCAPGly-mSc3 | This study | Lab plasmid |
| pEM791-4xCAPGly-mEGFP | This study | Lab plasmid |
| pEM791-4xCAPGly-mS3 | This study | Lab plasmid |
| pET15b-6×His-4xCAPGly-mEGFP | This study | Lab plasmid |
| pC1-superTagRFP-A1aY1 | This study | Lab plasmid |
| pEM791-superTagRFP-A1aY1 | This study | Lab plasmid |
| pET15b-6xHis-A1aY1-superTagRFP | This study | Lab plasmid |
| pCAGGS-internal PA-TubA1A-IRES-EGFP | This study | Lab plasmid |
| pCAGGS-internal PA-TubA1A(E254A)-IRES-EGFP | This study | Lab plasmid |
| pCAGGS-internal PA-TubA1A | This study | Lab plasmid |
| pCAGGS-internal PA-TubA1A(E254A) | This study | Lab plasmid |
| pCAGGS-mEGFP-TubA WT | This study | Lab plasmid |
| pCAGGS-mEGFP-TubA WT | This study | Lab plasmid |
| pPA-MAP2-mEGFP | This study | Lab plasmid |
| pPA-MAP7-mEGFP | This study | Lab plasmid |
| pN1-KIFC(1-560,G235A)-mEGFP | This study | Lab plasmid |
| pmEGFP-Tau | This study | Lab plasmid |
| pmEGFP-CAMSAP2 | This study | Lab plasmid |
| pmEGFP-CAMSAP3 | This study | Lab plasmid |
| pCAGGS-mEGFP-Tau | This study | Lab plasmid |
| pCAGGS- MAP2-mEGFP | This study | Lab plasmid |
| pCAGGS- mEGFP-CAMSAP2 | This study | Lab plasmid |
| pCAGGS- mEGFP-CAMSAP3 | This study | Lab plasmid |
| pFastBac1- KIF5C(1-560)-Halo-2xstrepII | This study | Lab plasmid |
| **Software and algorithms** | | |
| Fiji/ImageJ | Schneider et al. 2012 | https://fiji.sc/ |
| Fiji Lpx_bilevelThin plugin | Higaki et al. 2010 | |
| Fiji Ridge Detection plugin | Wagner et al. 2017 | |
| Fiji EzColocalization plugin | Stauffer et al. 2018 | |
| AIVIA | DRVision | |
| R version 4.3.1 | R Core Team | https://www.R-project.org/ |
| Bio3D | Grant et al. 2006 | |
| Prism version 10.4.1 | GraphPad Software | https://www.graphpad.com |
| Adobe Illustrator version 29.0 | Adobe | |
