## Supplemental File 1 for "Accessibility of the unstructured α-tubulin C-terminal tail is controlled by microtubule lattice conformation" (from another version of this preprint: version 2, 2025-12-19)

**Supplementary File 1. Primers used in cloning**

| | |
|---|---|
| CAPGLY_for1 | CTGCAGTCGACGGTACCGCGGCCCGGATGCTGAAACCCAGCGGGCTG |
| CAPGLY_rev1 | CACCTCCCGATCCACCACCGCCCCTCTGGAGTTTGTCAGCTTTGGTCTTTTC |
| CAPGLY_for2 | GTACCGCGGGCCCGGGATCCAATG CTGAAACCCAGCGGGCTG |
| CAPGLY_revGFP | GCCTGCACCTGAGTGTTTACTTTTACTTGTACAGCTC GTCCATGC |
| CAPGLY_revSc3 | CTGCACCTGAGTGTTTACTTTTAGGAGCCACCGGAGCC |
| CAPGLY_for3 | GCGGCAGCCA TATGCTCGAGCTGAAACCCAGCGGGCTGAAG |
| CAPGLY_rev3 | CTTTCGGGCTTTGTTAGCAGCCG TTACTTGTACAGCTCGTCCATGC |
| A1aY1_for1 | GATCTCGAGCTCAAGCTTCGGCAACAGTCAAGTTCAAATAC |
| A1aY1_rev1 | CGCGGTACCGTCGACTGCAGTTACTTCTTCTGCTTCTCC |
| A1aY1_for2 | TACCGCGGGCCCGGGATCCAATGGTGAGCAAGGGCGAG |
| A1aY1_rev2 | CTGCACCTGAGTGTTTACTTTTACTTCTTCTGCTTCTCCAGC |
| A1aY1_for3 | GCGGCAGCCATATGCTCGAGATGGCAACAGTCAAGTTCAAATAC |
| A1aY1_rev3 | CCATGGTACCCTTCTTCTGCTTCTCCAG |
| sTagRFP_for1 | GCAGAAGAAGGGTACCATGGTGAGCAAGGGCGAG |
| sTagRFP_rev1 | TCGGGCTTTGTTAGCAGCCGTTACTTGTACAGCTCGTCCATG |
| His-PA_for | TGCCGAAGATGATGTGGTGGGGGGAGGAGATGATTCCTTC |
| His-PA_rev | CCTGGCATGGCAACGCCAATGGTCTTGTCACTTGGCATC |
| IRES-GFP_removal_for | AGCGGCCGCAATTCACTCCTC |
| IRES-mEGFP_removal_rev | TTAGTATTCCTCTCCTTCTTC |
| mEGFP-TubA1A_for | CTCAAGCTTCGAATTCTGCAATGGTGAGCAAGGGCGAG |
| mEGFP-TubA1A_rev | AGGAGTGAATTGCGGCCGCTTTAGTATTCCTCTCCTTCTTCCTCAC |
| pCIG2_rev | TGCAGAATTCGAAGCTTGAGC |
| PA-mEGFP-N1_for1 | GGTGGAGGCGGTTCAGGC |
| PA-mEGFP-N1_rev1 | CACCACATCATCTTCGGCACC |
| MAP2_for | GTGCCGAAGATGATGTGGTGATGGCAGATGAACGGAAAG |
| MAP2_rev | CCGCCTGAACCGCCTCCACCCAAGCCCTGCTTAGCGAG |
| MmMAP7_for | GTGCCGAAGATGATGTGGTGGCGGAGCAGGGAGCTGGC |
| MmMAP7_rev | CCGCCTGAACCGCCTCCACCTATAACTTCTGCGGTCTGTTGTGTCTGCAC |
| PA-mEGFP-N1_for2 | GGCGTTGCCATGCCAGGTGCCGAAGATGATGTGGTGAGCAAGGGCGAGGA GCTGTTC |
| PA-mEGFP-N1_rev2 | CATGGTGGCGACCGGTGG |
| Kif5C_for | ATCCACCGGTCGCCACCATGGCGGATCCAGCCGAATGC |
| Kif5C_rev | GCACCTGGCATGGCAACGCCCTCGAGCGGATCCCGGGC |
| tau_for | GATCTCGAGCTCAAGCTTCGATGGCTGAGCCCCGCCAG |
| tau_rev | CGCGGTACCGTCGACTGCAGTCACAAACCCTGCTTGGCCAG |
| CAMSAP2_for | GATCTCGAGCTCAAGCTTCGATGGGGGATGCTGCAGACCCCA |
| CAMSAP2_rev | CGCGGTACCGTCGACTGCAGCTATGCCTTAGTGGGTAAAAGTTTTTTGG |
| CAMSAP3_for | GATCTCGAGCTCAAGCTTCGATGGAGATCAAGTCGCTGGACCA GTACGATTTCTCGCGG |
| CAMSAP3_rev | CGCGGTACCGTCGACTGCAGCTATTTGGGAGTGCCGCC |
| pCAGGS_for | AGCGGCCGCAATTCACTCCTC |
| pCAGGS_rev | AATTCGAAGCTTGAGCTCGAG |
| PA-mEGFP_MAP_for | TCGAGCTCAAGCTTCGAATTATGGGCGTTGCCATGCCA |
| PA-mEGFP_MAP_rev | AGGAGTGAATTGCGGCCGCTTTATCTAGATCCGGTGGATCCCG |
| PA-MAP2-mEGFP_rev | AGGAGTGAATTGCGGCCGCTTTACTTGTACAGCTCGTCCATGC |
